# The Reading Frame Surveillance Hypothesis

**DOI:** 10.1101/071985

**Authors:** John T. Gray

## Abstract

**Abbreviations:**

RFS: Reading Frame Surveillance
RdRP: RNA-dependent RNA Polymerase
frRNAs: Framing RNAs
LSU: Large Subunit
SSU: Small Subunit
tRF: Transfer RNA derived Fragment
nt: nucleotide

**Abstract:** An alternative model for protein translation is presented wherein ribosomes utilize a complementary RNA copy of protein coding sequences to monitor the progress of messenger RNAs during their translation to reduce the frequency of frameshifting errors. The synthesis of this ‘framing RNA’ is postulated to be catalyzed by the small subunit of the ribosome, in the decoding center, by excising and concatemerizing tRNA anticodons bound to each codon of the mRNA template. Various components of the model are supported by previous observations of tRNA mutants that impact ribosomal frameshifting, unique globin-coding RNAs in developing erythroblasts, and the epigenetic, intergenerational transfer of phenotypic traits via mammalian sperm RNA. Confirmation of the proposed translation mechanism is experimentally tractable and might significantly enhance our understanding of several fundamental biological processes.

## The Reading Frame Surveillance Hypothesis

The impetus for this alternative model of protein translation was a theoretical speculation as to how a ribosome might mechanistically prevent +1 frameshifting when translating an mRNA. Reading of mRNA by the ribosome entails direct binding of tRNA anti-codons to each codon of the message (Figure 1), and steric clashes between adjacent tRNAs should theoretically limit −1 frameshifting to some degree, as that event requires that two tRNAs be bound to the same base on the mRNA, or that the mRNA contain a short homopolymeric sequence that allows slippage of the P-site tRNA to make room for binding in the A-site after the frameshifting two-nucleotide translocation event, as has been described for extant translating ribosomes (Chamorro et al., 1992). Plus-one frameshifting, on the other hand, could occur without such steric inhibition whenever a four-nucleotide translocation allowed binding of the A-site tRNA in a +1-shifted position relative to the previous codon.

**Figure 1.**
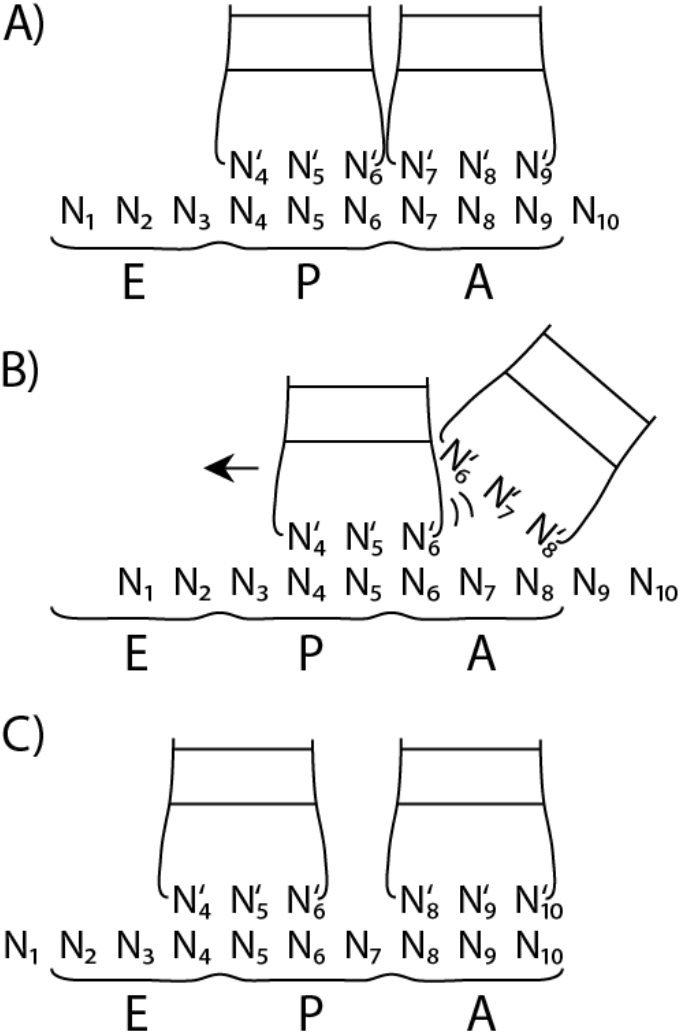
Steric Clashes Generated During −1 but Not +1 Frameshifting Positioning of A and P site tRNA anticodon stems are shown with (A) proper positioning, (B) −1 frameshifting resulting from a 2-nucleotide translocation event, without the necessary sliding of the P-site tRNA (arrow), and (C) +1 frameshifting resulting from a 4-nucleotide translocation event, with no steric clashes.

Prompted by the suggestion that genomic RNA replication in an ‘RNA World’ would cause primordial translation templates to be double stranded (Zenkin, 2012), I consider in this manuscript how and whether extant ribosomes might use a complementary RNA strand to measure the progress of a template RNA through the ribosome during translation to help prevent or respond to frameshifting events. This Reading Frame Surveillance (RFS) hypothesis posits that RNA annealed to translation templates could contain reading frame demarcating landmarks, (every third nucleotide or multiples thereof), and be sensed by the ribosome to detect or control the position of the mRNA relative to the reading frame being read. Figure 2 diagrams how in-frame (Figure 2A) and +1 frameshifted (Figure 2B) mRNA positioning could be sensed by the ribosome via two landmarks, nucleotide base modifications and RNA chain termini, on a complementary ‘framing’ RNA. Some aspects of the model remain undefined, such as whether the landmarks are sensed during each ribosomal translocation event to prevent frameshifting, or only periodically to facilitate regulatory responses to frameshifting events. One defining feature of the mechanism is the sensing of landmarks on the complementary framing RNA by a part of the ribosome complex at a fixed distance from the A, P, and E sites, to directly measure the position of the correct reading frame on the mRNA.

**Figure 2.**
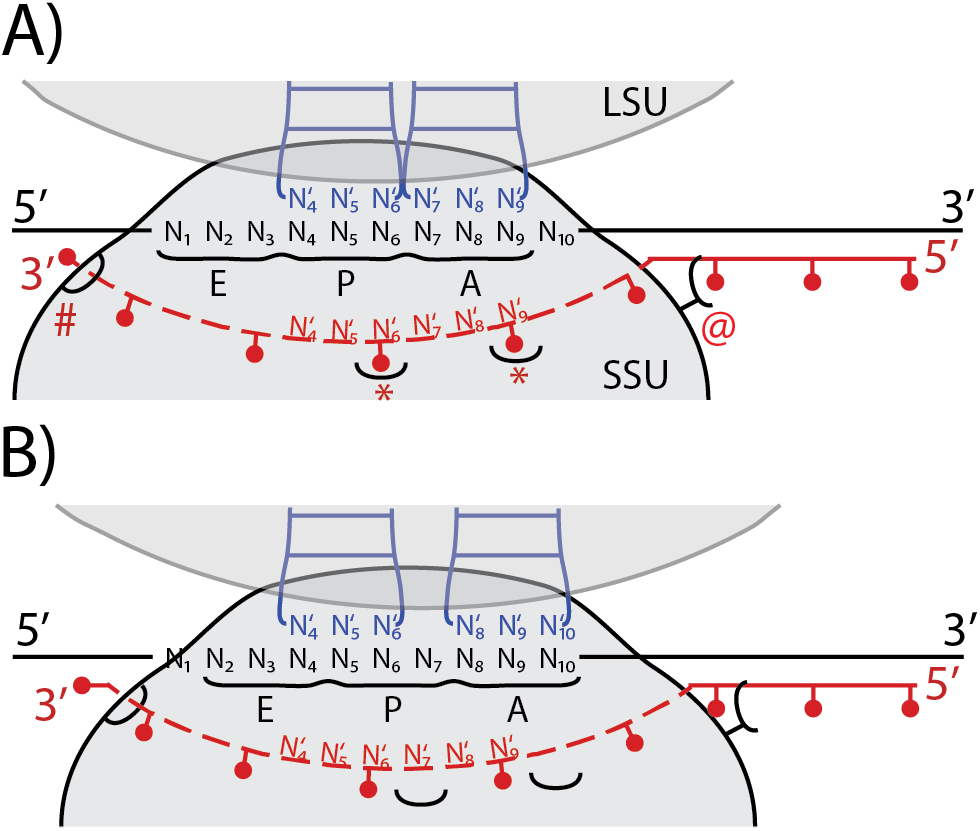
Reading Frame Surveillance Model for Protein Translation. A schematized view of the ribosomal small subunit (SSU) decoding center is shown, with the A, P, and E sites marked, along with multiple models for how framing RNA landmarks might be sensed by the ribosome. The template mRNA is shown as individual nucleotides in the decoding site (N_1-10_), and black lines externally. Anticodon stems of tRNAs bound by the ribosome are shown in blue, and a minimal framing RNA (frRNA) is shown in red, displaced from the mRNA template by the decoding center active site (dotted line). Red dots represent postulated modified nucleotides and termini on the frRNA, which could be sensed by the ribosome at several positions as the mRNA migrates across the decoding center active site. A) In-frame alignment, with nucleotides 4-6 in the P site, and 7-9 in the A site. Based on pairing of the frRNA to template sequences 3’ of the active site, the frRNA landmarks are positioned properly. LSU, large subunit; SSU, small subunit. ‘*’ indicates sensing of wobble nucleotide modifications in framing RNAs dissociated from but cognate for the codons in the A and P sites. ‘@’ indicates sensing of wobble modifications where the framing RNA is paired with the mRNA, here shown upstream of the active site. ‘#’ indicates sensing of framing RNA termini, in this case downstream of the active site. B) Frameshifted alignment, produced by a 4- nucleotide translocation which shifts the small subunit forward such that an incorrect tRNA can bind in the A-site in a new reading frame. This repositions the frRNA landmarks such that they are no longer in line with the wobble position of the A, P, and E sites.

Central to the question of whether such framing RNAs exist in cells is how they could possibly be synthesized and marked to support reading frame detection. Synthesis of an RNA complementary to each codon in a pulsatile, three-nucleotides-at-a-time cycle could allow that RNA to be marked by base modification at a consistent position opposite each codon. Such landmarks could be sensed as the corresponding codons are translated (as indicated by ‘*’ in Figure 2), or outside the ribosome active site when the framing RNA is paired to the template (e.g., ‘@’ in Figure 2). Additionally, if the enzyme performing the synthesis was not fully processive, then the termini of multiple, shorter framing RNAs could be positioned at a consistent position with respect to the reading frame and allow periodic sensing of the frame by detection of RNA chain termini (‘#’ in Figure 2). In another synthesis model, an already double stranded RNA is marked or cleaved at regularly spaced intervals during a pioneer round of translation such that the landmarks are positioned properly for later translation events. This pioneer translation model, however, requires yet another model to explain how pioneer translation would be differentiated to achieve lower rates of frameshifting than would occur normally without the presence of framing RNAs. Thus, the ‘sets of three’ synthesis model is currently preferred, as it appears to provide the most straightforward mechanism whereby the framing RNAs could be marked at a uniform position within each codon, and also potentially restrict the positions of termini on framing RNAs.

The well understood role for the small subunit of the ribosome in translation initiation and mRNA decoding, combined with recent observations of tRNA fragments in cells (Lee et al., 2009), prompted further speculation that the small subunit of extant ribosomes might utilize the ancient decoding center active site to perform the postulated framing RNA synthesis, similar to previously suggested mechanisms for primordial translation (Poole et al., 1999). Figure 3A details a hypothetical series of transesterification reactions between tRNAs bound in the SSU active site that effectively result in RNA dependent RNA polymerization (see details in figure legend). Such concatenation of tRNA anticodon loops could create framing RNAs with covalent modifications opposite the wobble position of each codon that function as framesensing landmarks during later translation of the same message. A model for how this framing RNA synthesis by the SSU might integrate with protein translation in cells is shown in Figure 3B, with framing RNA synthesis occurring as part of the process of translation initiation.

The core features of this Reading Frame Surveillance hypothesis, namely that ribosomes generate an RNA copy of translation templates from tRNA anticodons and use those molecules as guides during translation to maintain proper reading frame, is speculative and significantly challenges well accepted models for extant protein synthesis. Yet published data supports the existence of an RNA dependent RNA polymerase (RdRP) in vertebrate cells (Kapranov et al., 2010; Tseng and Lai, 2009; Volloch et al., 1996), even though this activity has never been attributed to any protein enzyme after decades of effort. Although the absence of robust sequence data documenting cellular RNAs complementary to coding sequences appears to contradict the tenets of the model, just because such framing RNAs have not been described does not mean they do not exist. In fact, some support for the existence of RNAs that could function as framing RNAs not only exists, as detailed below, but also provides a rationale for why they have escaped detection to date.

## Nucleotide Base Modifications in tRNA and the Mechanism of +1 Frameshifting

Nucleotides 34, the wobble position in the anticodon loop, and 37, the first nucleotide after the anticodon, are two of the most highly modified bases in all RNAs, and are typically modified in nearly all tRNAs in all domains of life (Grosjean et al., 2010; Jackman and Alfonzo, 2013). Extensive research has documented how the modification of these residues impacts protein synthesis, (reviewed in (Agris et al., 2007; Atkins and Bjork, 2009)), notably in their ability to modulate the frequency of +1 frameshifting in both prokaryotic and eukaryotic systems. In *E. coli*, modifications of nucleotides in and adjacent to the anticodon stem significantly enhance the maintenance of reading frame by preventing +1 but not −1 frameshifting in multiple tRNA species (Urbonavicius et al., 2001; Urbonavicius et al., 2003). Current postulated mechanisms whereby a lack of base modification enhances frameshifting are varied and sometimes not clear, although they frequently invoke a failure of unmodified tRNAs to compete with tRNAs cognate for +1-frameshifted codons in the ribosomal active sites (Atkins and Bjork, 2009), an idea supported by the observation that the unmodified tRNAs do not contribute amino acids to the growing polypeptide chain when frameshifting occurs (Qian et al., 1998).

While the binding of a tRNA to a frameshifted mRNA codon in the ribosomal A site is clearly a key step for ribosomal frameshifting, it can only occur after a non-canonical 4 nt translocation event positions the codon in the A-site. The Reading Frame Surveillance model proposes that framing RNAs could facilitate control of translocation to prevent such an event, and the covalent modifications at position 34 of tRNA anticodon loops, after incorporation into framing RNAs, may instead function as the proposed landmarks that ribosomes use to maintain that control. Some covalent modifications at position 34 are bulky yet do not interfere with the base pairing face of the nucleotide (see Figure 4 for chemical structures of some modifications observed at position 34 of tRNAs), making them suitable for this role. Thus, mutation of the enzymes responsible for those modifications may result in framing RNAs lacking landmarks opposite the codons where frameshifting has been shown to occur, (or in fact at other critical positions along a coding sequence), and this lack of framing RNA landmarks could facilitate aberrant 4 nucleotide translocation events that allow recognition of a +1-frameshifted codon by an alternative tRNA.

**Figure 4.**
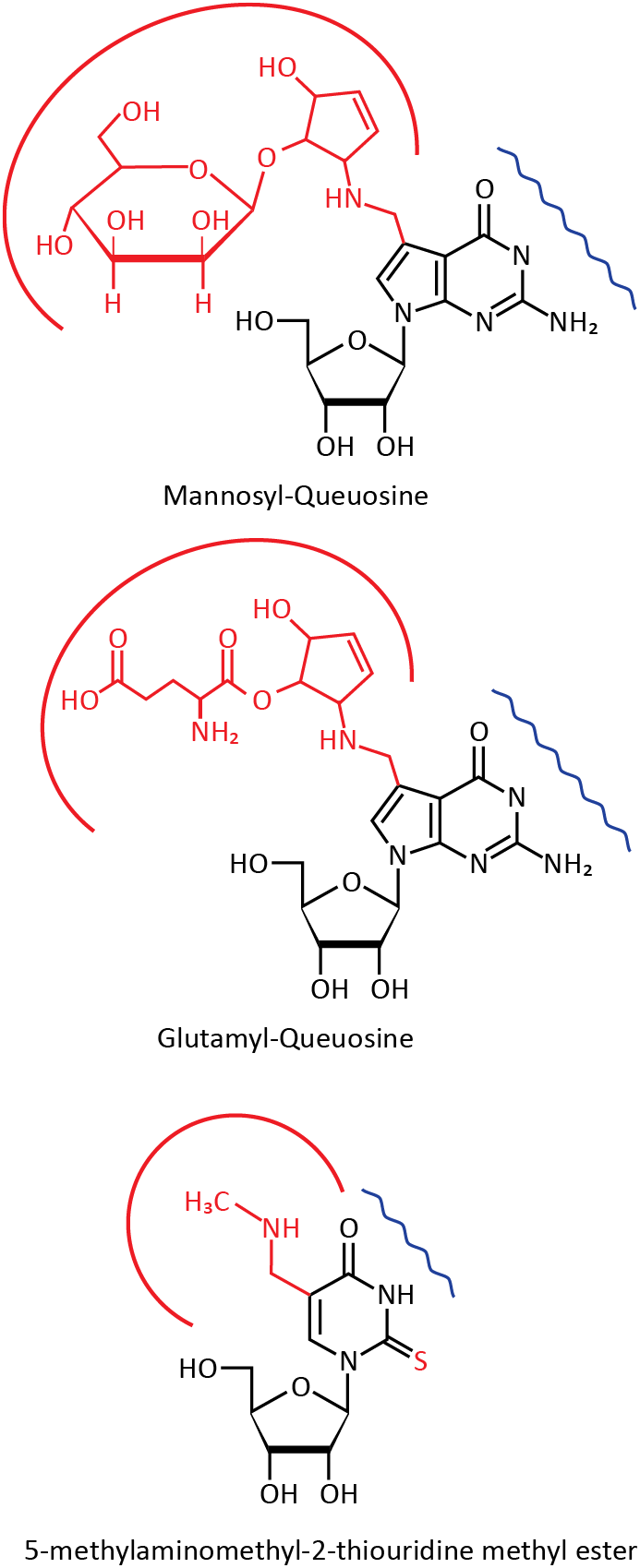
Structures of Modified Bases Observed at Position 34 of tRNAs Three nucleotide base modifications are shown, with the non-canonical atoms shown in red. Mannosyl-queuosine is a modified form of guanosine found in eukaryotes, and glutamyl-queuosine is found in prokaryotes. 5-methylaminomethyl-2-thiouridine methyl ester is found in both prokaryotes and eukaryotes. The base pairing face of each modified nucleotide is marked with a blue wavy line, while bulky groups that could serve as framing RNA landmarks are partially circled in red. When paired to mRNA, the bulky groups would be expected to protrude into the major groove of the helix. Structures are as presented at http://modomics.genesilico.pl (Machnicka et al., 2013).

The observed promotion of +1 frameshifting by two additional forms of altered tRNA anticodon loop structures is similarly consistent with the RFS model. Both insertion of an extra nucleotide into the anticodon loop and mutations in enzymes responsible for 1- methylguanosine modification of position 37 (which blocks base pairing by the modified base) have been modeled to allow base pairing of 4 anticodon loop nucleotides to the mRNA at positions where those altered tRNAs facilitate +1 frameshifting (Bjork et al., 1989; Riddle and Carbon, 1973). These observations originally prompted a ‘yardstick model’ whereby pairing of tRNA anticodon loops somehow measures the proper length of ribosomal translocation, but these models were difficult to reconcile with the previously mentioned observation that the mutant tRNAs are not utilized by the ribosome during translation events that shift frames (Qian et al., 1998). By the RFS model, these frameshifting events can be explained by the aberrant incorporation of 4 nucleotides into framing RNA, which shifts the spacing of landmarks and facilitates frameshifting during translation by promoting 4 nucleotide translocation events.

Although some frameshift-promoting tRNA alterations (e.g., those distal to the anticodon loop) are not obviously explained by the RFS model, the above well studied examples do support the model, and warrant its inclusion as a possible mechanistic framework for interpreting the consequences of tRNA mutations. Conversely, the conservation of base modifications at positions 34 and 37 of the anticodon loop in all domains of life, the lack of a uniform, accepted mechanistic rule as to why they are necessary, and the abundant genetic evidence that sequences of and nucleotide base modifications in anticodon loops can alter the frequency of +1 frameshifting could be interpreted as tentative and subjective evidence supporting the RFS model.

## Globin mRNA in Developing Erythroblasts

Characterization of globin protein coding RNAs in developing erythroblasts has established that RNAs like framing RNAs do exist, at least in vertebrate cells, and have structures consistent with synthesis from tRNA anticodon loops. Twenty years ago, globin mRNAs were first proposed to be copied and amplified by an unidentified cellular RdRP, (Volloch et al., 1996), but the idea has failed to gain broad acceptance in the absence of definitive molecular cloning and sequencing of the amplified RNA sequences. In the original Volloch et al. publication, globin antisense RNA molecules were identified by strand specific northern blotting, but could not be molecularly cloned as cDNAs and sequenced. More recent experiments have confirmed that these antisense RNAs are used as a template in erythroblasts during generation of large quantities of sense strand RNAs that were both mapped by RNase protection assays to confirm they possessed the globin gene sequence, and translated in vitro to show they coded for globin protein (Rits et al., 2016).

Notably, however, these globin end-product RNAs (comparable in abundance to ribosomal RNAs themselves) were shown to possess modified nucleotides throughout their length (550600 nucleotides) which prevented their conversion to cDNA and characterization by conventional methods. Reverse transcriptase enzymes are inherent to all molecular methods for cloning and sequencing of RNA, and yet the typical enzymes employed cannot copy tRNA anticodon loops because of the nucleotide base modifications, (see, for example, (Qian et al., 1998) for an example). I therefore suggest that these globin coding RNAs could in fact be synthesized by the ribosome SSU from tRNA anticodon loops via something like the three-at-a-time mechanism proposed in Figure 3, and accordingly are covalently modified at every third nucleotide, preventing cloning and sequencing by conventional methods. Thus, although these RNAs are not antisense to the coding strand, and likely result from an additional round of synthesis by the ribosome to meet the unique needs of globin protein synthesis in developing erythroblasts, their abundance and extensive modification demonstrates that the enzymatic function at the heart of the RFS model not only exists in extant cells but can be extraordinarily robust.

**Figure 3.**
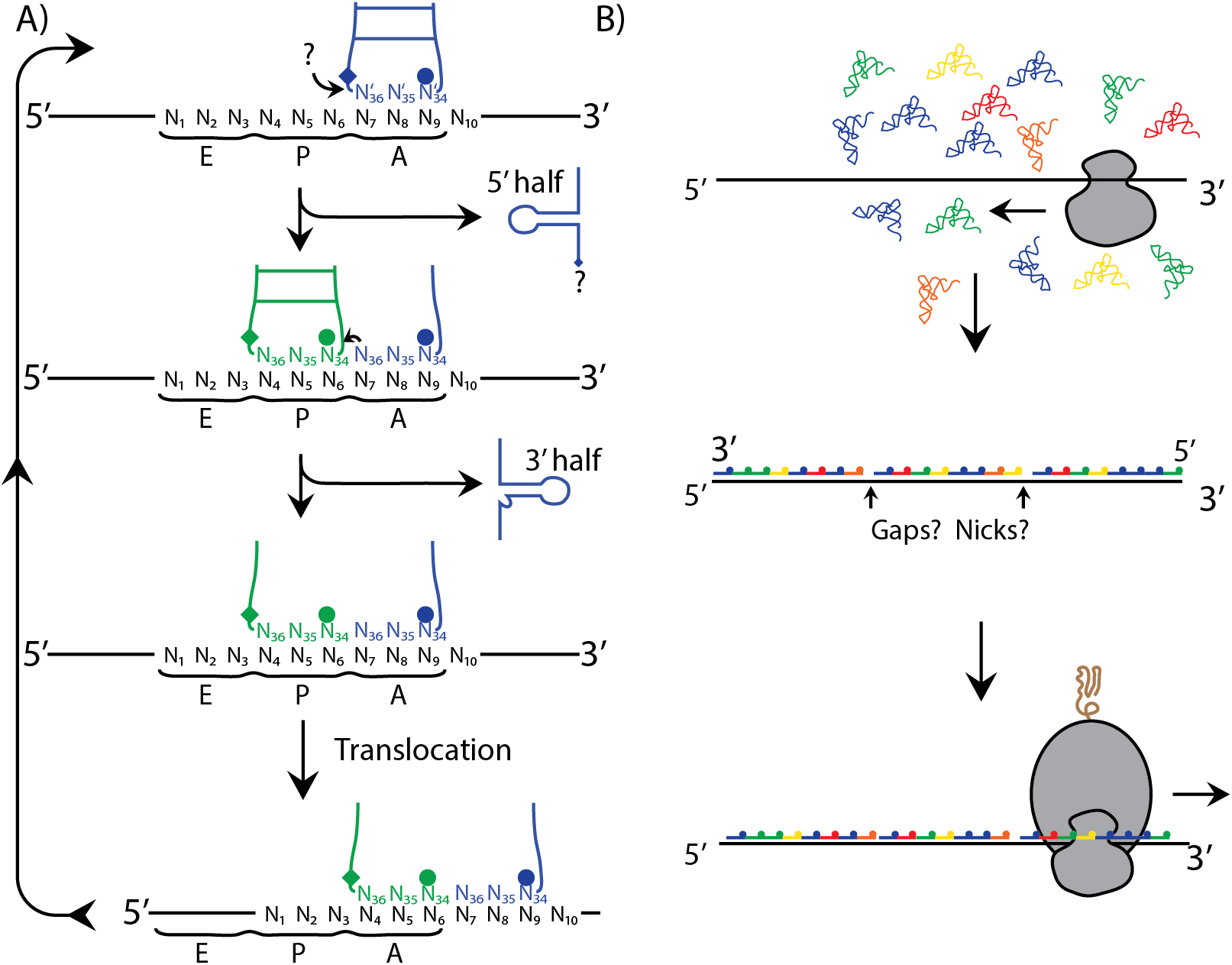
Framing RNA Synthesis by the Ribosomal Small Subunit Decoding Center. A) A hypothetical model for a series of transesterification events between tRNAs bound in the A and P sites is presented, although similar steps might just as easily occur in the P and E sites, and independent cleavage and ligation events are also consistent with the model. In this figure the tRNA nucleotide numbering follows convention for tRNAs, with nucleotides (nts) 34-36 encompassing the anticodon, and nucleotide 34 as the wobble position. Position 34 modifications are indicated by opaque circles, position 37 as diamonds. An unknown hydroxyl, perhaps a 3’ OH from a free nucleotide, (indicated by ‘?’) initiates the process by attacking the phosphate on the 5’ side of the modified residue 37, joining to and releasing the 5’ half of the first tRNA. The remaining 3’ hydroxyl is then free to attack 5’ of the modified residue 34 of the next incoming tRNA, releasing the 3’ half of that tRNA. Translocation allows repetition of the process to elongate the growing chain in a template dependent manner in the 5’ to 3’ direction, although cleavage and ligation reactions need not bear such directionality. B) The hypothesized action of the SSU to create framing RNAs is illustrated as part of protein translation initiation. Each tRNA is colored to denote different anticodon sequences. Synthesis is speculatively modeled as 5’ to 3’ based on the transesterification chemistry presented in panel A. Regardless, incorporation of anticodons into a growing polymer would position modified residues at the 5’ end of each triplet on the framing RNA (the wobble position, indicated by dots), which would be recognized during subsequent translation by the complete ribosome.

## Transfer RNA Fragments and the Epigenetic Regulation of Gene Expression

The last argument in support of the RFS model is less evidence than intriguing speculation, which is that framing RNAs might be novel candidates for RNA-mediated transfer of epigenetic information, as has been demonstrated for many small RNAs bound to Argonaute family proteins (Aravin et al., 2008; Tabara et al., 1999; Yamanaka et al., 2013), some of which are complementary to protein coding sequences (Conine et al., 2013; Seth et al., 2013; Shirayama et al., 2012). To explore this possibility, consider the non-Mendelian, epigenetic inheritance of the phenotypic traits of fur color (Kiani et al., 2013; Rassoulzadegan et al., 2006) and glucose tolerance (Carone et al., 2010; Chen et al., 2016; Sharma et al., 2016). In these murine systems, it is believed that RNA transmitted via sperm from male sires during fertilization is capable of affecting gene expression in developed progeny, yet the specific RNA responsible for this epigenetic inheritance has not been established (see (Rando, 2016) for a review of intergenerational epigenetic transfer in sperm).

Sperm contain abundant quantities of tRNA fragments (tRFs), (Peng et al., 2012), which have recently gained interest as important signals of cellular stress (Fu et al., 2009; Lee et al., 2009; Thompson et al., 2008), and these sperm tRFs were investigated as potential mediators of the above mentioned epigenetic phenotypes (Chen et al., 2016; Kiani et al., 2013; Sharma et al., 2016). Unfortunately, although total RNA purified from sperm was shown to confer the epigenetic phenotype to progeny mice when injected into embryos, and in one case this activity was determined to reside in the 30-40 nt size fraction containing tRFs, synthetic versions of these tRFs failed to elicit the same response (Chen et al., 2016; Kiani et al., 2013). Framing RNA synthesis, as modeled in Figure 3A, is expected to generate similar 5' and 3' tRFs, and so the generation of tRFs in the epididymal caput during sperm development (Sharma et al., 2016) could in fact be evidence of a developmentally regulated burst of framing RNA synthesis. This interpretation suggests that we should consider whether framing RNAs are the critical epigenetic factor mediating these phenotypes. This hypothesis is supported by the lack of evidence for significant alterations in gene expression from tRFs, as mentioned, (Chen et al., 2016; Kiani et al., 2013), the lack of significant information content in tRF molecules relative to the number of genes that have been shown to be affected by epigenetic factors in sperm (>400, according to (Carone et al., 2010)), and the alternatively large data capacity available to framing RNAs (essentially complementing any translated sequence).

Relevant to this question is the observation that mutation of *Dnmt2*, which encodes a methylase that modifies tRNA outside the anticodon loop, was shown in mice to prevent the epigenetic transmission of fur color paramutants (Kiani et al., 2013). Extensive data supports the role of base modifications in tRNA structure stabilization (Helm, 2006), and modification by Dnmt2 and another tRNA methylase, (NSun2) have been shown to protect tRNAs from cleavage into tRFs by angiogenin, a stress activated ribonuclease. Mutations in *Dnmt2* and *NSun2* increase, rather than decrease, cellular stress (Blanco et al., 2014; Schaefer et al., 2010). Although the failure of *Dnmt2* mutant mice to transmit fur color paramutations could be interpreted to mean that covalently modified tRFs are the critical mediators of that process (Chen et al., 2016), it is equally supportive of epigenetic transmission requiring properly structured tRNAs to facilitate framing RNA synthesis, which are themselves the communicating factor. In this view, failures in the RFS pathway cause cellular stress by decreasing translational accuracy and creating misfolded proteins, and the cleavage of misfolded tRNAs is one of many possible responses that can facilitate regeneration of the translation apparatus to restore cell health. This perspective does not necessarily contradict the proposed direct influence by tRFs on stress response pathways (Thompson and Parker, 2009) and endogenous retroviral LTR driven transcription (Sharma et al., 2016).

## Experimental Validation of the Reading Frame Surveillance Model

Although framing RNAs seem to have generally escaped detection so far, growing interest in how RNA modifications influence cell physiology has led to significant improvements in the analysis of modified RNAs using a variety of techniques (Clark et al., 2016; Cozen et al., 2015; Mishima et al., 2015; Zhang et al., 2016; Zheng et al., 2015). Specifically, treatment of tRNAs with demethylases and use of a novel thermostable reverse transcriptase can significantly enhance the ability to read through tRNA modifications (Cozen et al., 2015; Zheng et al., 2015), and implementation of some or all of these techniques with strand specific library preparation methods will allow discrimination of reads from those derived from coding mRNAs. Such an analysis, followed by northern blot characterization of select framing RNAs, should allow confirmation of the existence of framing RNAs in cells, and if present determine whether they are short, as are many epigenetic factors (Chen et al., 2016; Joshua-Tor and Hannon, 2011), or the full length of coding sequences, as were found for globin antisense RNAs in erythroblasts (Volloch et al., 1996).

Genetic approaches, in particular those that helped characterize the impact of tRNA mutations on frameshifting frequencies (Atkins and Bjork, 2009), may also provide evidence for the RFS mechanism. The vast quantities of data generated with tRNA anticodon stem mutants and tRNA modification enzyme mutants using translation reporter templates could be evaluated with the RFS model in mind, to perhaps confirm the model or even identify a particular genetic context in which it is most strongly active. For direct testing of the model, it may be possible to construct tRNA mutant classes that segregate the framing RNA synthesis vs. decoding activities of the ribosome, for example by preventing charging by tRNA synthetases while allowing normal incorporation into framing RNAs. Such mutants, when provided into cells carrying frameshifting anticodon mutant tRNAs, might revert their phenotype, which would support the inclusion of those anticodons into framing RNAs. Such experiments might also provide an indication as to whether framing RNA landmarks are sensed concurrently with translocation of the matching mRNA codon into the decoding center active site, (i.e., when framing RNA is displaced from the A, P, and E sites, marked as ‘*’ in Figure 2), or whether sensing can occur upstream or downstream when the framing RNA is paired with the mRNA, ('@' and ‘#’ in Figure 2), to provide a more generalized monitoring of reading frame maintenance across a transcript.

Ideally, the RFS model can be confirmed by demonstrating framing RNA synthesis with ribosomal preparations *in vitro*, and it is not clear that this has ever been directly tested. *In silico* experiments were performed to address the question of whether tRNA anticodon loops positioned in the A and P sites could ligate with each other, but in translating ribosomes the sites were too far apart (Noller, 2012). It is intriguing to consider whether the canonical L-shaped structure of tRNAs and the large subunit peptidyl transferase center active site hold anticodon loops in a position that prevents their concatenation during protein synthesis, while during translation initiation in the absence of the LSU an alternative alignment juxtaposes them appropriately for framing RNA synthesis. It may thus be worthwhile to evaluate structures of ribosomal small subunits incubated with mRNA templates and tRNAs to see a more relevant structure. The most stringent test for catalytic activity of a ribozyme is to use purely synthetic RNA components, which also facilitates incorporation of easily detected radioactive or chemical labels. Or, purified ribosomes from cell types postulated to have RdRP activity, such as developing erythroblasts (Volloch et al., 1996), HDV infected cells (Tseng and Lai, 2009), or epididymal epithelia (Sharma et al., 2016) might be more likely to show activity concordant with increased quantities of tRNA fragments.

## Broader Implications of the RFS Model

Generally, the possibility that cells could contain RNA molecules complementary to any coding sequence, and that these RNAs have to date largely escaped detection, should be considered as relevant to many biological questions. Framing RNA pairing with mRNA transcripts could play a role in the selection for GC-rich codons in coding exons (Amit et al., 2012; Louie et al., 2003), the particular position and spacing requirements for stop codons in nonsense mediated decay pathways (Popp and Maquat, 2014), the impact of translatability on exon inclusion choices (Miriami et al., 2004), and the epigenetic communication of gene silencing (Guzzardo et al., 2013; Xu et al., 2015; Yamanaka et al., 2013) and alternative splicing patterns (Yokota et al., 2006). Likewise, the RFS pathway in general may prove to be relevant to the initiation of cellular stress, (as discussed above), the generation of antigenic peptides for presentation by the Major Histocompatibility Complex I (Apcher et al., 2015; Dolan et al., 2011), and the regulation of expression of certain alternative reading frame tumor suppressors (Sherr, 2012).

Finally, the presence of an RdRP enzyme in vertebrates has been debated for decades (Downey et al., 1973; Kapranov et al., 2010; Tseng and Lai, 2009; Volloch et al., 1996; Zong et al., 2009), and it is hoped that the experiments suggested in this manuscript will bring some resolution to the debate. Although RdRP ribozymes are commonly thought to have existed in RNA-based primordial living systems (Crick, 1968), the suggestion that such an enzyme is present in extant cells does appear to have been considered. The first discovery of catalytic RNA thirty-five years ago (Kruger et al., 1982) initiated the search for extant ribozymes, and 10 years later the idea that the large subunit ribosomal RNA directly catalyzes the peptidyl transferase activity at the heart of protein translation was largely accepted (Noller et al., 1992). The hypothesis presented here attempts to extend those findings, by providing a mechanistic justification for a unique form of RdRP activity as part of protein translation, and showing how the small subunit of the ribosome and tRNA anticodon base modifications are structurally suited to performing that function. As the small and large subunit ribosomal RNAs are two of the most highly conserved genetic sequences in all life on earth, and protein translation is uniformly essential to all extant life, the suggested pairing of RNA replicase and peptidyl transferase activities in extant small and large subunit ribozymes has an appealing symmetry, and if confirmed, may significantly enhance our understanding of the origins of life.

## Acknowledgements

The author has been generously supported by Audentes Therapeutics, Inc., the American Lebanese Syrian Associated Charities, and the Assisi Foundation of Memphis. The author is currently a salaried employee of Audentes Therapeutics, Inc., owns equity interest in that company, and receives royalty payments resulting from the licensing of gene transfer technology by St. Jude Children's Research Hospital.

